# Memorability of Images Positively Influences Associative Memory Retention

**DOI:** 10.1101/2025.09.03.674110

**Authors:** Ye Wang, Meixuan Li, Lei Yang, Lingfang Tan, Tong Li, Hongjiang Xiao, Yaling Deng, Yuan Zhang

## Abstract

Enhancing memory function is essential for daily living and cognitive health, particularly amid population aging and cognitive decline. However, current methods for memory enhancement often require specialized interventions or effortful practice. Here, we present evidence that the intrinsic memorability of images can serve as a simple, scalable, and involuntary strategy for improving associative memory. We studied its effects on associative memory in adults of distinct ages, memory retention in cognitively impaired older adults, and vocabulary learning in foreign language learners. Our results show that highly memorable images significantly enhance the recall of associated words, especially when the cue image and target word are semantically unrelated. Notably, these effects persist for at least one week and are robust across age groups and cognitive status. In a foreign language vocabulary task, pairing words with memorable images led to either improved recall accuracy or reduced learning time. These findings highlight that, in addition to being an intrinsic memory-enhancing property, image memorability also serves as a general facilitator of associative memory, via a method that is easy to adopt, requires minimal conscious effort, and can benefit the general public, particularly learners and cognitive impairment patients in educational and clinical settings.

## Introduction

Given the pivotal role of memory enhancement in education, aging, and clinical interventions, improving memory performance has long been a central objective in cognitive psychology and neuroscience. Although a variety of methods have been developed to enhance memory, each approach faces major limitations that hinder widespread adoption. Techniques such as transcranial direct current stimulation (tDCS) ^1, 2^ and transcranial magnetic stimulation (TMS) ^3, 4^ require specialized equipment and are not easily accessible. Pharmacological interventions can cause unwanted side effects^5,6,7^, and cognitive training typically demands intensive, sustained effort, which limits its practicality and scalability ^8, 9,10,11, 12^. These limitations are especially salient in educational contexts, among older adults, and in populations with cognitive impairments. As such, identifying a memory enhancement strategy that is natural, easy to implement, and free from side effects has remained a major challenge of cognitive psychology and neuroscience.

Recent studies have shown that certain visual stimuli, particularly highly memorable images, can involuntarily influence memory consolidation ^13, 14^. This involuntary effect is an intrinsic property of the image itself^15,16^, independent of a viewer’s attention^17^, motivation^18^, or personal preferences^15^. Notably, the memory advantage conferred by these highly memorable images can last from a week^19^ to several years^20^. However, prior researches have focused almost exclusively on memory for the images themselves^16,21,22^. Given that effective memory in everyday life often depends on linking visual stimuli to other relevant information, understanding the impact of image memorability on associative memory is especially important. Yet whether this benefit extends beyond the images themselves remains unclear.

To address this gap, we conducted a series of experiments to test whether highly memorable images could enhance associative memory retention beyond the image itself. Across distinct age groups and cognitive status, as well as in a second-language learning context, we found that pairing information with highly memorable images consistently improved memory retention, especially when the images and associative items were semantically unrelated. Notably, these benefits persisted for at least one week. Our findings suggest that image memorability acts not only as an intrinsic memory-enhancing property, but also as a general facilitator of associative memory, serving as a cognitive anchor and opening new directions for memory enhancement research.

## Highly memorable images enhance associative memory under low semantic relatedness

To determine whether image memorability can function as a memory aid for recalling associated information, we conducted a preregistered image-word associative memory experiment (Experiment 1; https://osf.io/u98zs). Previous research has shown that greater semantic relatedness between items typically leads to stronger associative memory^23^. To account for this, we created two semantic relatedness conditions: high and low. Within each, we further manipulated image memorability (high vs. low), resulting in four experimental conditions: high memorability image + high semantic relatedness word (HM+HS), high memorability image + low semantic relatedness word (HM+LS), low memorability image + high semantic relatedness word (LM+HS), and low memorability image + low semantic relatedness word (LM+LS), as illustrated in **Fig. 1a**. See Supplementary Information Section 1.1 for details in stimuli selection.

**Figure 1.**
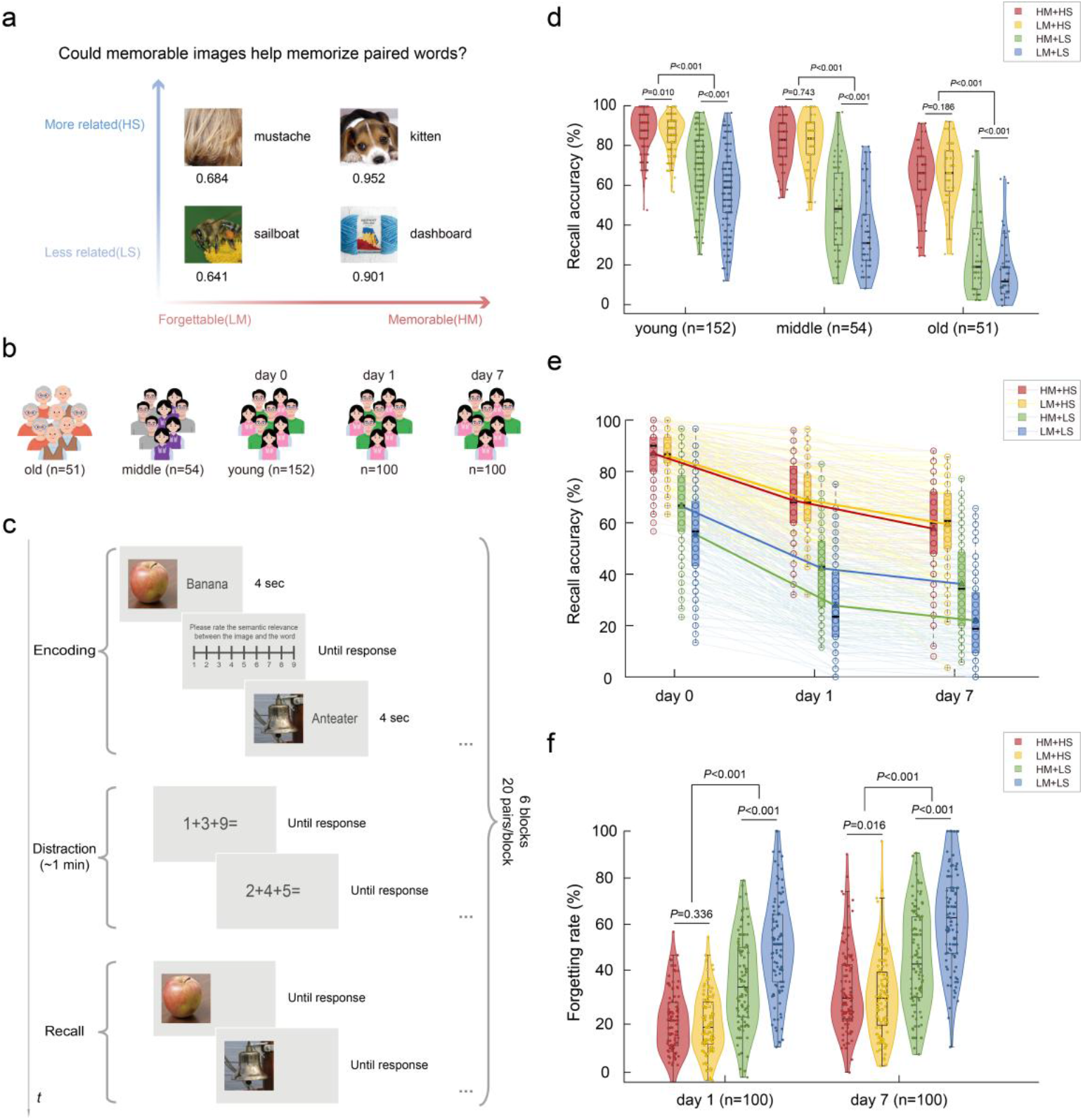
Highly memorable images enhance associative memory when semantic relatedness is low. **a**, Schematic of the research question: Can highly memorable (HM) images serve as effective cues to facilitate recall of paired words? The numbers below the pictures represent the memorability scores. **b**, Participant groups and follow-up design for Experiment 1. Young adults (18–34 years, *n* = 152), middle-aged adults (35–54 years, *n* = 54), and older adults (≥55 years, *n* = 51) were recruited to complete the initial test; a subset of young adults (*n* = 100) participated follow-up test at 1- and 7-days post-learning. **c**, Overview of the experimental procedure: Each session included an encoding phase (4 seconds per image-word pair, with semantic relatedness ratings on a 1–9 Likert scale), a 1-minute distraction task (single-digit arithmetic), and a recall phase (image cue only, participant recalls the paired word). The main task comprised six blocks of 20 pairs, with all four experimental conditions evenly distributed. **d**, Recall accuracy by age group and experimental condition. The highest recall was observed in the high semantic relatedness (HS) condition for both HM and low-memorability (LM) images. Under low semantic relatedness (LS), HM images led to significantly higher recall than LM images. **e**, The memory enhancement effects of HM and HS are stable on the test day (day 0), 1 day later, and 7 days later. The young subjects showed a certain degree of forgetting of the image-word pairs in the experiment over time. The triangles in the box plot represent the mean, and the short black line represents the median. **f**, Forgetting rates over time by condition. The lowest forgetting rates were found in the HS condition regardless of memorability. Under LS conditions, HM images produced lower forgetting rates than LM images, with this benefit sustained for at least one week.

To examine the effects of image memorability on associative memory among distinct age groups and over time, we recruited participants from three age groups: young adults (18–34 years), middle-aged adults (35–54 years), and older adults (55 years and above). In addition, a subset of young participants completed follow-up tests one day and seven days after the initial test (**Fig. 1b**). The initial session consisted of encoding, distraction, and recall (**Fig. 1c**), while follow-up tests assessed recall only.

To determine whether recall accuracy differed across groups and experimental conditions, we conducted a 2 (memorability: high, low) × 2 (semantic relatedness: high, low) × 3 (age group: young, middle-aged, older) mixed-design analysis of variance (ANOVA), with age group as a between-subjects factor. We found a significant main effect of memorability (*F*(1, 254) = 111.074, *P* < 0.001, *η*^2^ = 0.304), with participants recalling more words paired with high-memorability (HM) images (*M* = 63.066%, *SE* = 0.970%) than with low-memorability (LM) images (*M* = 57.755%, *SE* = 0.948%; mean difference = 5.311%, 95% confidence interval (CI) = [4.318%, 6.303%], *P* < 0.001). There was also a significant main effect of semantic relatedness (*F*(1, 254) = 1383.690, *P* < 0.001, *η*^2^ = 0.845): image-word pairs with high semantic relatedness (HS) were recalled more accurately (*M* = 78.277%, *SE* = 0.743%) than those with low semantic relatedness (LS) (*M* = 42.544%, *SE* = 1.274%; mean difference = 35.733%, 95% CI = [33.841%, 37.625%], *P* < 0.001). The main effect of age group was also significant (*F*(2, 254) = 119.310, *P* < 0.001, *η*^2^ = 0.484): young adults (*M* = 75.492%, *SE* = 1.065%) performed significantly better than middle-aged adults (*M* = 62.707%, *SE* = 1.787%; mean difference = 12.785%, 95% CI = [7.771%, 17.798%], *P* < 0.001), who, in turn, outperformed older adults (*M* = 43.032%, *SE* = 1.839%; mean difference = 19.675%, 95% CI = [13.495%, 25.854%], *P* < 0.001).

A significant interaction was observed between memorability and semantic relatedness (*F*(1, 254) = 126.278, *P* < 0.001, *η*^2^ = 0.332). In the HS condition, recall accuracy did not differ significantly between HM and LM images (*F*(1, 256) = 1.421, *P* = 0.234, *η*^2^ = 0.006; mean difference = 0.692%, 95% CI = [-0.451%, 1.836%]). In the LS condition, however, the recall was significantly higher for HM images than LM images (*F*(1, 256) = 286.85, *P* < 0.001, *η*^2^ = 0.528; mean difference = 10.924%, 95% CI = [9.653%, 12.194%]). This pattern was consistent across all age groups (see Supplementary Information Section 2.1 for details).

These results indicate that, across all age groups, the HS condition yielded the highest memory performance, regardless of whether the image was high or low in memorability. In the LS condition, pairing words with HM images led to significantly better recall than pairing with LM images, as shown in **Fig. 1d**.

For young participants who completed both the 1-day and 7-day follow-up sessions, we conducted a 2 (memorability: high, low) × 2 (semantic relatedness: high, low) × 2 (time: 1 day, 7 days) repeated-measures ANOVA to assess differences in forgetting rates across conditions. There was a significant main effect of time (*F*(1, 99) = 119.125, *P* < 0.001, *η*^2^ = 0.546): forgetting rates were higher at 7 days (*M* = 43.981%, *SE* = 1.622%) than at 1 day (*M* = 32.625%, *SE* = 1.360%; mean difference = 11.356%, 95% CI = [9.292%, 13.421%], *P* < 0.001). Memorability also had a significant main effect (*F*(1, 99) = 63.258, *P* < 0.001, *η*^2^ = 0.390), with HM images showing lower forgetting rates (*M* = 34.849%, *SE* = 1.439%) than LM images (*M* = 41.758%, *SE* = 1.499%; mean difference = -6.909%, 95% CI = [-8.632%, -5.185%], *P* < 0.001). The main effect of semantic relatedness was significant as well (*F*(1, 99) = 383.550, *P* < 0.001, *η*^2^ = 0.795): HS pairs had significantly lower forgetting rates (*M* = 26.707%, *SE* = 1.252%) than LS pairs (*M* = 49.899%, *SE* = 1.753%; mean difference = -23.193%, 95% CI = [-25.542%, -20.843%], *P* < 0.001). A significant three-way interaction was observed (*F*(1, 99) = 4.318, *P* = 0.040, *η*^2^ = 0.042).

Further simple-simple effects analyses revealed that, under the LS condition, HM images had significantly lower forgetting rates than LM images both at the 1-day delay (*F*(1, 99) = 92.151, *P* < 0.001, *η*^2^ = 0.482; mean difference = -14.945%, 95% CI = [-18.034%, -11.856%]) and 7-day delay (*F*(1, 99) = 115.040, *P* < 0.001, *η*^2^ = 0.537; mean difference = -16.612%, 95% CI = [-19.685%, -13.539%]). In the HS condition, there was no significant difference in forgetting rates between HM and LM images at 1-day delay (*F*(1, 99) = 0.933, *P* = 0.336, *η*^2^ = 0.009; mean difference = 1.097%, 95% CI = [-1.157%, 3.352%]), but after 7 days, HM images showed significantly higher forgetting rates than LM images (*F*(1, 99) = 5.983, *P* = 0.016, *η*^2^ = 0.057; mean difference = 2.825%, 95% CI = [0.533%, 5.116%]).

These results indicate that associative memory for image-word pairs declined across all conditions as the retention interval increased. Nonetheless, the memory enhancement effects of both high memorability (HM) and high semantic relatedness (HS) persisted for at least seven days. Specifically, the lowest forgetting rates were observed in the HS condition, regardless of image memorability. Under the LS condition, HM images led to significantly lower forgetting rates compared to LM images (**Fig. 1e-f**).

### Highly memorable images enhance associative memory in elderly individuals with cognitive impairment

To examine whether the memory benefits of highly memorable (HM) images apply to individuals with cognitive impairment and whether they help reduce forgetting over time in older adults, we conducted Experiment 2 with 100 elderly participants (age: 55– 96 years, *M* = 73.76, *SD* = 10.14; 24 males, 76 females). All participants completed the Montreal Cognitive Assessment (MoCA)^24^ and were classified as cognitively normal (MoCA score ≥ 26, *n* = 52; age: 55–94, *M* = 72.60, *SD* = 10.49; 11 males, 41 females; MoCA score: 26–30, *M* = 27.49, *SD* = 1.29) or cognitively impaired (MoCA score < 26, *n* = 48; age: 55–96, *M* = 75.02, *SD* = 9.70; 13 males, 35 females; MoCA score: 15– 25, *M* = 21.94, *SD* = 2.56). Among these, 82 participants completed the 1-day follow-up, and 78 completed both the 1-day and 7-day follow-up sessions (**Fig. 2a**).

**Figure 2.**
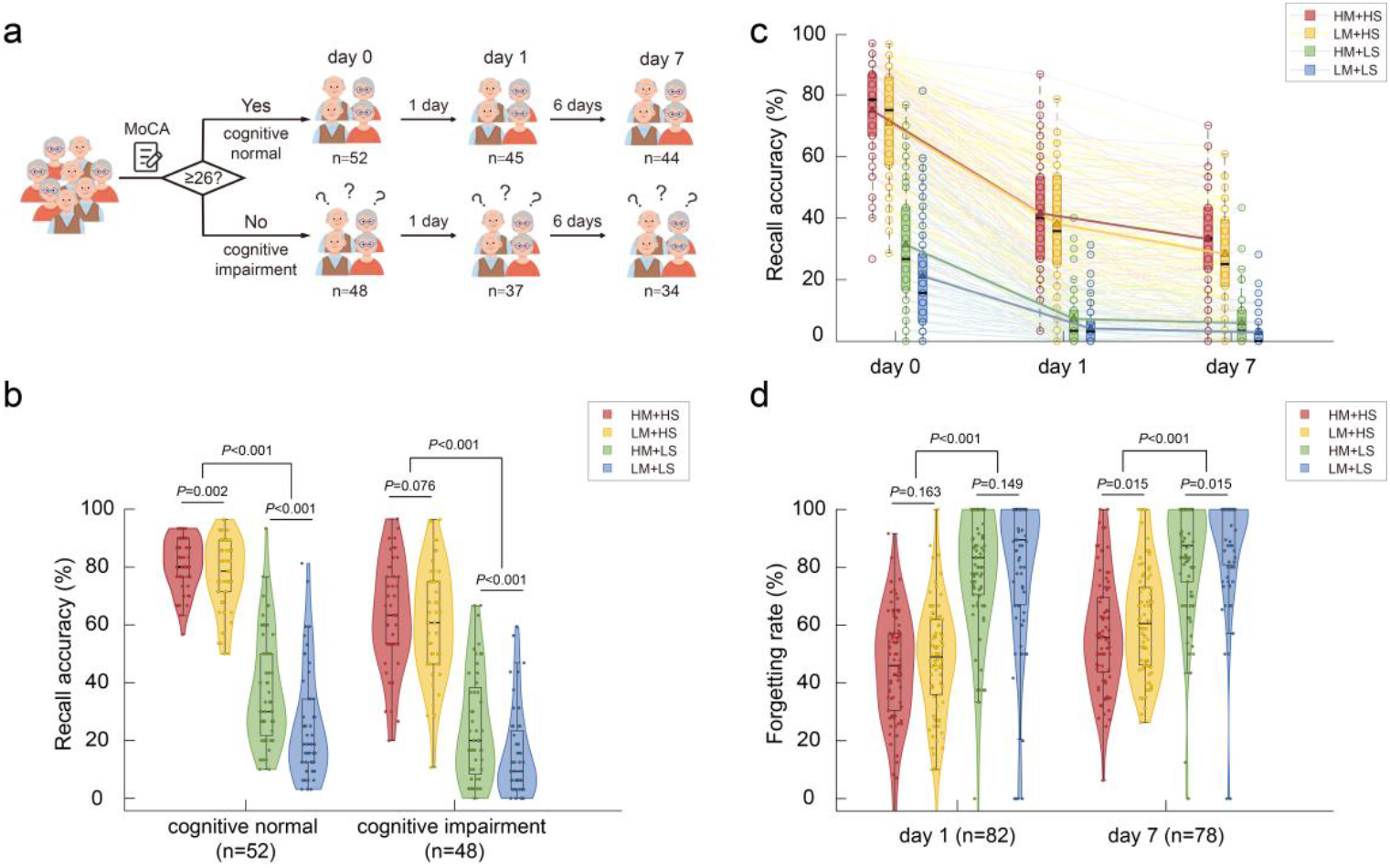
Highly memorable images enhance associative memory in older adults with cognitive impairment. **a**, Schematic of Experiment 2 design and participant grouping. A total of 100 older adults (ages 55–96) participated and completed the Montreal Cognitive Assessment (MoCA). Based on their scores, 52 were classified as cognitively normal (MoCA score ≥ 26) and 48 as cognitively impaired (MoCA score < 26). Of these, 82 completed a 1-day follow-up, and 78 completed both 1-day and 7-day follow-ups. **b**, In both cognitively normal and cognitively impaired groups, pairing words with highly memorable (HM) images significantly improved recall accuracy compared to low-memorability (LM) images, especially under low semantic relatedness (LS) conditions. **c**, Recall accuracy declined over time for all elderly participants, with greater forgetting at longer intervals after learning. The triangles in the box plot represent the mean, and the short black line represents the median. **d**, The lowest forgetting rates were found in the low semantic relatedness (HS) condition regardless of memorability. In the 7-day follow-up, HM images resulted in lower forgetting rates regardless of semantic relatedness.

Since Experiment 2 used different stimulus materials from Experiment 1 (see Supplementary Information Section 1.2 for details), we tested the generalizability of Experiment 1’s findings by comparing recall rates between adults with normal cognitive aging and those with cognitive impairment across four experimental conditions. We conducted a 2 (memorability: high, low) × 2 (semantic relatedness: high, low) × 2 (group: cognitively normal, cognitively impaired) mixed-design ANOVA, with the group as the between-subjects factor. There was a significant main effect of memorability (*F*(1, 98) = 129.193, *P* < 0.001, *η*^2^ = 0.569): recall accuracy was higher for HM images (*M* = 51.535%, *SE* = 1.502%) than for LM images (*M* = 44.306%, *SE* = 1.465%), with a mean difference of 7.229% (95% CI = [5.967%, 8.492%], *P* < 0.001). Semantic relatedness also showed a significant main effect (*F*(1, 98) = 1067.928, *P* < 0.001, *η*^2^ = 0.916): HS pairs (*M* = 70.454%, *SE* = 1.425%) were recalled more accurately than LS pairs (*M* = 25.387%, *SE* = 1.766%), with a mean difference of 45.066% (95% CI = [42.330%, 47.803%], *P* < 0.001). The main effect of group was significant as well (*F*(1, 98) = 24.148, *P* < 0.001, *η*^2^ = 0.198), with cognitively normal participants (*M* = 55.042%, *SE* = 2.008%) recalling more words than the cognitively impaired group (*M* = 40.799%, *SE* = 2.090%; mean difference = 14.242%, 95% CI = [8.491%, 19.994%], *P* < 0.001). The interaction effects of memorability and other factors are reported in Supplementary Information Section 2.2. Notably, the memory advantage of HM over LM images was greater under LS conditions than under HS conditions (see Supplementary Information Section 2.3 for details).

We further analyzed whether the interaction between semantic relatedness and memorability was evident in both cognitively normal and cognitively impaired older adults. Among cognitively normal participants, recall accuracy was significantly higher for HM than for LM images in both the HS (*F*(1, 51) = 10.838, *P* = 0.002, *η*^2^ = 0.175; mean difference = 3.956%, 95% CI = [1.544%, 6.368%]) and LS conditions (*F*(1, 51) = 107.826, *P* < 0.001, *η*^2^ = 0.679; mean difference = 12.416%, 95% CI = [10.015%, 14.816%]). In the cognitively impaired group, there was no significant difference between HM and LM images under the HS condition (*F*(1, 47) = 3.284, *P* = 0.076, *η*^2^ = 0.065; mean difference = 2.629%, 95% CI = [-0.290%, 5.548%]); however, under the LS condition, recall was significantly higher for HM than for LM images (*F*(1, 47) = 49.804, *P* < 0.001, *η*^2^ = 0.514; mean difference = 9.916%, 95% CI = [7.090%, 12.743%]).

These results indicate that the main effect of memorability and its interaction with semantic relatedness were observed in both cognitively normal and cognitively impaired groups, with consistent trends. Under the LS condition, HM images led to higher recall of paired words than LM images in both groups (**Fig. 2b**).

We further examined differences in forgetting rates across conditions for all older adult participants in Experiment 2. As shown in **Fig. 2c**, memory for image-word pairs declined over time, with longer intervals leading to greater forgetting. After a 1-day delay, there was no significant difference in forgetting rates between HM and LM images under either the HS (*F*(1, 81) = 1.980, *P* = 0.163, *η*^2^ = 0.024; mean difference = -2.448%, 95% CI = [-5.910%, 1.014%]) or LS (*F*(1, 77) = 2.120, *P* = 0.149, *η*^2^ = 0.027; mean difference = -5.805%, 95% CI = [-13.745%, 2.135%]) conditions. In contrast, after 7 days, forgetting rates were significantly lower for HM images than for LM images under both HS (*F*(1, 77) = 6.177, *P* = 0.015, *η*^2^ = 0.074; mean difference = - 4.141%, 95% CI = [-7.458%, -0.823%]) and LS (*F*(1, 73) = 6.194, *P* = 0.015, *η*^2^ = 0.078; mean difference = -5.284%, 95% CI = [-9.515%, -1.052%]) conditions. These results show that the memory-preserving effect of HM images persisted for at least one week (**Fig. 2d**). Similar time-related patterns of forgetting were observed for both adults with normal cognitive aging and those with cognitive impairment (see Supplementary Information Section 2.4 for details).

### Highly memorable images enhance foreign language vocabulary learning efficiency

Pairing foreign vocabulary words with images that represent their meanings (for example, associating the word “library” with a real-world photo of a library) is a common learning strategy. However, because learners generally lack prior knowledge of the foreign words, the semantic link between the words and their paired images is often weak. This raises the question: Can highly memorable (HM) images enhance the efficiency of foreign vocabulary learning in such contexts, as suggested by our findings in Experiments 1 and 2, where HM images improved recall under low semantic relatedness (HM+LS > LM+LS; **Fig. 3a**)?

**Figure 3.**
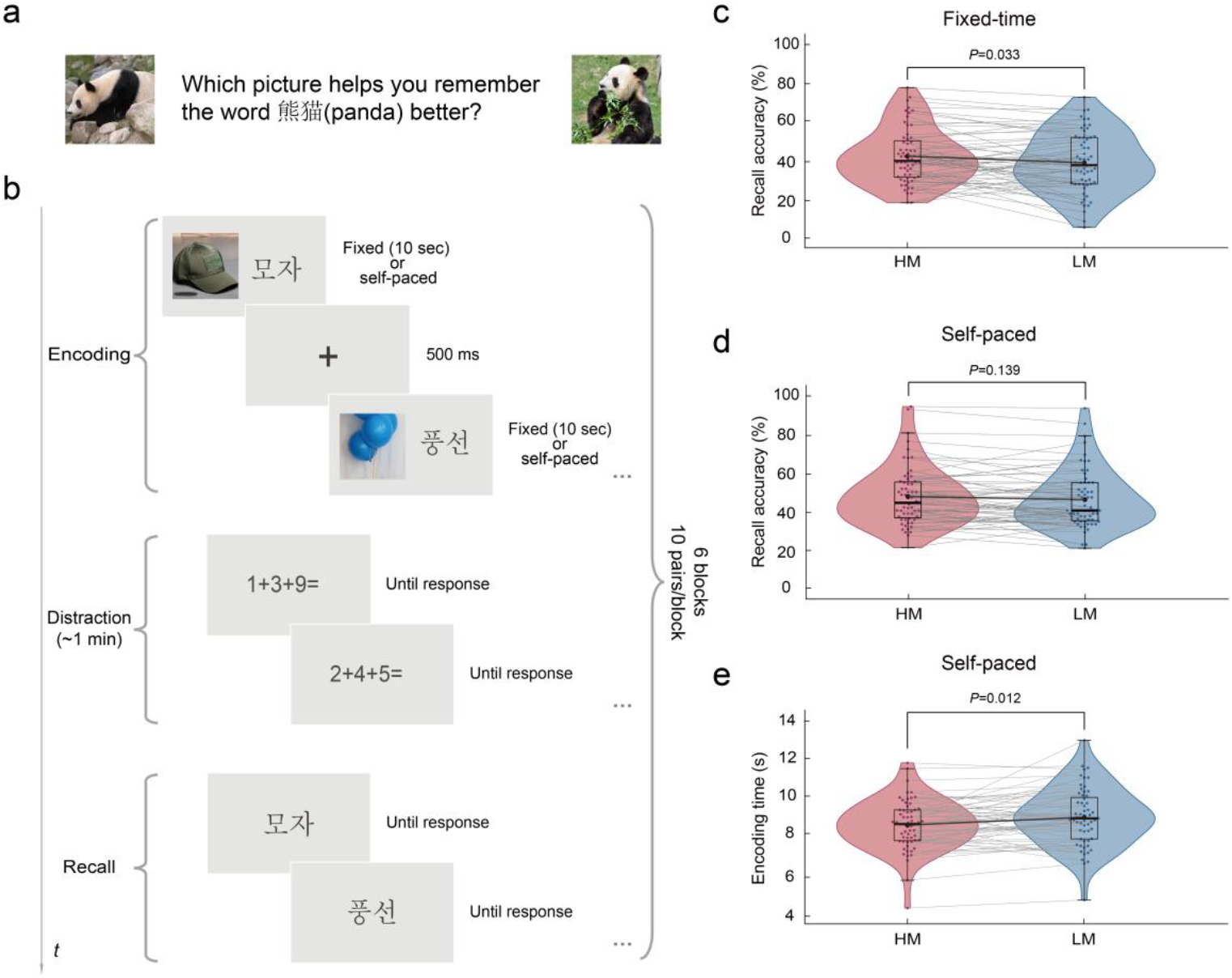
Highly memorable images improve foreign language vocabulary learning efficiency. **a**, Schematic of the experimental question: Does pairing words with highly memorable (HM) images improve the efficiency of foreign language vocabulary learning compared to low-memorability (LM) images? **b**, Experimental design for Experiment 3. Participants learned foreign vocabulary by pairing each word with either an HM or LM image in two conditions: fixed presentation time (10 seconds per pair) and self-paced presentation time (participants controlled when to advance). Each block comprised encoding, a 1-minute distractor task, and a recall phase in which participants provided the meaning for each word. The main task included six blocks of 10 pairs, with HM and LM images evenly distributed. **c**, Recall accuracy under fixed-time condition. Words paired with HM images showed significantly higher recall accuracy than those paired with LM images. **d**, Recall accuracy under self-paced condition. Recall accuracy did not differ significantly between words paired with HM and LM images. **e**, Encoding time under self-paced condition. Participants spent significantly less time encoding words paired with HM images than those paired with LM images, indicating greater learning efficiency.

To answer this, we conducted Experiment 3 with Chinese college students, using two learning paradigms: a fixed-time condition (each image-word pair was presented for 10 seconds) and a self-paced condition (participants controlled the presentation duration, advancing when they believed they had learned the pair). Both tasks included encoding, distraction, and recall phases (see Supplementary Information Section 1.3 for details; **Fig. 3b**). Memory efficiency was assessed both by recall accuracy and encoding time.

Paired-sample t-tests revealed that, in the fixed-time condition, HM images led to significantly higher recall accuracy for words (*M* = 42.835%, *SE* = 1.815%) than LM images (*M* = 39.496%, *SE* = 1.978%; *t*(59) = 2.184, two-sided *P* = 0.033, *d* = 0.282; mean difference = 3.339%, 95% CI = [0.280%, 6.397%]), as shown in **Fig. 3c**. In the self-paced condition, there was no significant difference in recall accuracy between HM (*M* = 48.414%, *SE* = 1.945%) and LM images (*M* = 46.786%, *SE* = 1.933%; *t*(59) = 1.500, two-sided *P* = 0.139, *d* = 0.194; mean difference = 1.628%, 95% CI = [-0.544%, 3.799%]), as shown in **Fig. 3d**. However, analysis of encoding times showed that HM images (*M* = 8.682 sec, *SE* = 0.166) significantly reduced the time needed to memorize words compared to LM images (*M* = 9.079 sec, *SE* = 0.191; *t*(59) = -2.592, two-sided *P* = 0.012, *d* = -0.335; mean difference = -0.398, 95% CI = [-0.705, -0.091]), as shown in **Fig. 3e**.

## Discussion

Our data demonstrate that highly memorable images serve as effective enhancers of associative memory. This enhancement effect persists for at least one week, generalizes across age groups, as well as individuals with cognitive impairment, as shown in Experiments 1 and 2. Additionally, we show that this benefit extends to foreign language vocabulary learning (Experiment 3), illustrating the generalizability of memorability-based facilitation across ages and learning contexts. By establishing these consistent effects across diverse populations and tasks, our findings deepen the current understanding of image memorability, suggesting that it is not merely a stimulus-intrinsic property but also a key factor facilitating memory integration and consolidation.

We provide converging evidences that image memorability can function as an involuntary, implicit and temporally stable memory aid, in contrast to traditional approaches ^25,26,27^ that require deliberate cognitive strategies or interventions. This involuntary enhancement is notably advantageous in contexts where active learning or sustained attention is challenging. These behavioral effects may arise from the brain’s inherent preference for processing memorable visual content, which could spontaneously trigger deeper encoding processes without deliberate cognitive effort^14,28^. To better understand the neural mechanisms underlying these behavioral effects, we propose that the medial temporal lobe (MTL), which is central to associative memory formation^29^, plays a key role in this process. Although neuroimaging studies have reported enhanced MTL activation for highly memorable stimuli ^30, 31^, single-unit recordings in humans have not shown increased neural firing rates for these images^32^. This discrepancy suggests that the stronger activation observed in neuroimaging studies is more likely driven by neural oscillations rather than by increased firing rates of individual neurons^33^. Indeed, neural oscillations, such as hippocampal ripples, are rhythmic patterns of neural activity that have been linked to episodic memory consolidation^34^. Recent research further shows that hippocampal ripple rates can predict image memorability during free recall^35^ and are thought to facilitate associative binding across stimulus-specific neurons^36^. Building on these findings, we hypothesize that highly memorable images may evoke increased ripple activity in the hippocampus during encoding, thereby promoting stronger associative memory traces. Testing this hypothesis will require future studies that combine intracranial recordings in humans with computational modeling, which could reveal the neural mechanisms that support ripple-mediated memory enhancement and connect them directly to the considerable behavioral effects we observed.

Because memorability-based strategies can involuntarily enhance associative memory, they are particularly well suited for practical applications in clinical, educational, and communication settings. In clinical contexts, highly memorable images have the potential to provide a non-invasive and scalable option for memory rehabilitation in individuals with mild cognitive impairment or early-stage Alzheimer’s disease. In education, incorporating memorable visual cues into teaching materials can help students retain new information more effectively, particularly in areas such as foreign language learning or abstract concept mastery. In communication applications such as marketing, journalism, or multimedia design, leveraging memorability can increase message retention and emotional engagement for broad audiences with the potential to create substantial commercial value. Altogether, these insights reveal memorability’s potential not only to transform how we strengthen and preserve memory, but also to empower individuals to learn, connect, and recover in ways that are more accessible, meaningful, and enduring, bringing us closer to a future where memory can enrich every aspect of life.

## Methods

Here, we provide an overview of participant recruitment, experimental procedures, and data preprocessing for Experiments 1, 2, and 3. Except for the follow-up tests in Experiment 1, all experimental stimuli were presented using PsychoPy (v2024.1.3), and all participants completed the experiments on-site in a quiet environment. The follow-up tests for Experiment 1 were conducted online using Wenjuanxing (wjx.cn), a survey platform functionally equivalent to Qualtrics or SurveyMonkey. All statistical analyses were conducted using SPSS 22.0 unless otherwise specified. All studies were approved by the Ethics Committee of the Communication University of China (Approval No. XSLL20241021-1), and informed consent was obtained from all participants.

### Experiment 1

#### Participants

To investigate the effects of image memorability on associative memory across age groups and over time, we recruited participants through advertisements in three age categories: young adults (18–34 years), middle-aged adults (35–54 years), and older adults (55 years and above). A total of 152 young adults (aged 18–21 years, *M* = 19.32, *SD* = 0.74; 33 males, 119 females), 54 middle-aged adults (35–54 years, *M* = 44.69, *SD* = 6.21; 23 males, 31 females), and 51 older adults (55–89 years, *M* = 66.39, *SD* = 9.58; 20 males, 31 females) participated in the initial test. All participants were native Chinese speakers. Of these, 100 young adults also completed follow-up tests at 1 and 7 days after the initial session. Participants received either course credit or a stipend of ¥30 (US$4.18).

#### Experimental design

The initial test consisted of three phases: encoding, distraction, and recall. During encoding, each image-word pair was presented for 4 seconds, after which participants rated the semantic relatedness of the pair on a 9-point Likert scale (1 = not related at all, 9 = highly related). A new image-word pair was shown following each rating. After all pairs had been presented, participants completed a 1-minute distraction task, which involved solving simple addition problems with three single-digit numbers. In the recall phase, participants were shown the cue image and asked to recall the associated word. Responses were either typed directly by the participant or, if the participant was unfamiliar with typing, spoken aloud and entered by the experimenter.

The session included both a practice phase and a formal experiment. The formal experiment comprised six blocks, each containing 20 image-word pairs, with the four stimulus conditions evenly distributed within each block. For each participant, both the order of blocks and the sequence of stimuli within each block were randomized. Before beginning the formal experiment, all participants completed a practice phase that followed the same procedure, but included only five image-word pairs and a 15-second distraction task. The entire session lasted approximately 40 minutes.

Follow-up tests were administered one and seven days after the initial test. Only the images shown during the initial test were presented, and participants were asked to recall the associated words. The order of image presentation was randomized. Each follow-up session took approximately 20 minutes.

#### Data Preprocessing

Due to factors such as language context, individual life experience, and the polysemy of words, semantic similarity ratings derived from word embeddings may not always align with participants’ subjective similarity judgments^37^. These subjective judgments may also vary across distinct age groups. Therefore, we collected similarity ratings for each concept pair from participants in each age group and used k-means clustering (implemented in Python v3.11.5 with scikit-learn v1.6.1) to assign each pair to either the high semantic similarity (HS) or low semantic similarity (LS) category for subsequent analyses. The resulting number of stimuli per condition in each age group is shown below:

|  | HM+HS | LM+HS | HM+LS | LM+LS |
| --- | --- | --- | --- | --- |
| **Young** | 25 | 28 | 35 | 32 |
| **Middle** | 24 | 25 | 36 | 35 |
| **Old** | 24 | 27 | 36 | 33 |

Three research assistants, blind to the experimental conditions, independently encoded participants’ responses. For partially correct responses (e.g., recalling “stool” for “footstool”), a response was considered correct if at least two of the three assistants agreed it constituted a partial recall.

### Experiment 2

#### Participants

A total of 100 older adults participated in Experiment 2. Participants received ¥30 (US$4.18) for completing the initial test, ¥60 (US$8.36) for completing both the initial and 1-day follow-up tests, and ¥100 (US$13.93) for completing all three tests (the initial test, 1-day, and 7-day follow-up tests).

#### Experimental design

As in Experiment 1, the initial test included encoding, distraction, and recall phases. To accommodate the cognitive abilities of older adults, we reduced the task difficulty by increasing the number of blocks and decreasing the number of image-word pairs per block. The formal experiment consisted of 15 blocks, each containing eight image-word pairs, with the four stimulus conditions evenly distributed within each block. Each distraction task lasted 30 seconds. For each participant, both the order of blocks and the sequence of stimuli within each block were randomized. Before the formal experiment, all participants completed a practice phase with three image-word pairs and a 15-second distraction task, following the same procedure as the formal session. The entire initial session lasted approximately 40 minutes.

Follow-up sessions were conducted one and seven days after the initial test. Only images from the initial test were presented in a randomized order, and participants were asked to recall the associated words. Each follow-up session lasted approximately 20 minutes.

#### Data Preprocessing

As in Experiment 1, subjective similarity ratings provided by participants sometimes differed from estimates derived from word embeddings.

Therefore, we obtained subjective similarity ratings for each concept pair and used k-means clustering (implemented in Python v3.11.5 with scikit-learn v1.6.1) to categorize each pair as either high semantic similarity (HS) or low semantic similarity (LS) for subsequent analyses. After reclassification, the numbers of pairs in each condition were as follows: HM+HS (30), LM+HS (28), HM+LS (30), and LM+LS (32). Response coding and statistical analyses were conducted as in Experiment 1.

### Experiment 3

#### Participants

We recruited 162 native Chinese-speaking undergraduate and graduate students (aged 18–25 years, *M* = 19.55, *SD* = 1.12; 35 males, 127 females) to participate in Experiment 3. Participants received either course credit or a stipend of ¥30 (US$4.18). All participants reported no familiarity with Korean.

#### Procedure

To assess the impact of image memorability on foreign language vocabulary learning, each Korean word was paired with both a high-memorability (HM) and a low-memorability (LM) image. These pairings were distributed across two experimental groups (A and B). Half of the participants (*n* = 79) were randomly assigned to Group A, and the other half (*n* = 80) to Group B. Each group was presented with 60 image-word pairs: 30 words paired with HM images and 30 with LM images. The pairing was counterbalanced such that words paired with HM images in Group A were paired with LM images in Group B, and vice versa. This design enabled within-item comparison of recall accuracy across memorability conditions between groups.

The efficiency of foreign language vocabulary learning is reflected not only in higher recall accuracy but also in shorter encoding time. Therefore, we designed two experimental paradigms: a fixed-time experiment, in which image-word pairs were shown for 10 seconds each, and a self-paced experiment, in which participants controlled the duration of each image-word pair and pressed a key to advance once they believed they had memorized the pair.

The fixed-time task consisted of three phases: encoding, distraction, and recall. During encoding, each image-word pair was shown for 10 seconds, followed by a 0.5-second cross-fixation. After all pairs had been presented, participants completed a 1-minute distraction task involving single-digit addition problems. In the recall phase, participants saw only the Korean word and were asked to recall the meaning represented by the paired image.

The main experiment consisted of six blocks, each containing 10 image-word pairs. The two conditions (HM and LM images) were evenly distributed within each block. For each participant, both the order of the blocks and the sequence of stimuli within each block were randomized. A brief practice phase preceded the main experiment, using three image-word pairs and a shortened 15-second distraction task to familiarize participants with the procedure. Participants completed the entire experiment in approximately 45 minutes.

The procedure of the self-paced experiment was identical to that of the fixed-time experiment, except that participants controlled the duration of each image-word pair and pressed a key to advance once they believed they had memorized the pair. Because some participants took part in both experiments, distinct sets of Korean words were used for each to avoid overlap. Each word was paired with both a high-memorability (HM) and a low-memorability (LM) image, distributed across two experimental groups (X and Y). Half of the participants (*n* = 80) were randomly assigned to Group X, and the other half (*n* = 80) to Group Y. Each group viewed 60 words, with 30 words paired with HM images and 30 with LM images. The pairings were counterbalanced such that words paired with HM images in Group × were paired with LM images in Group Y, and vice versa, enabling within-item comparison of recall accuracy across memorability conditions.

#### Response Coding

Three research assistants, blinded to the experimental conditions and hypotheses, independently encoded participants’ responses. A response was marked as correct if it reflected the meaning of the associated image. At least two out of three assistants had to agree for a response to be considered correct. Participants with recall accuracy below the chance level (10%) were excluded from the analysis. After these exclusions, the final sample sizes were: Group A (*n* = 68), Group B (*n* = 63), Group × (*n* = 73), and Group Y (*n* = 63).

To ensure the validity of the encoding time data, trials with encoding time shorter than 100 ms or more than three standard deviations above the mean were excluded, as such values likely reflected inattentive or anomalously delayed responses. In total, 534 trials were excluded, resulting in 8,160 valid trials for encoding time analysis.

## Data availability

The datasets generated during and/or analyzed during the current study are available from the corresponding author on reasonable request.

## Acknowledgements

This work was supported by the Fundamental Research Funds for the Central Universities (Grant Nos. CUC24SG008, CUC25SG006)

## Author contributions

Y.W., Y.D. and Y.Z. conceived of the research; Y.W., M.L., L.Y., L.T. and Y.D. designed the experiments; Y.W., M.L., L.T. and T.L. conducted the experiments; Y.W. and Y.D. analyzed the data; Y.W., L.Y. and Y.D. wrote the paper, with input from M.L., L.T., T.L. and Y.Z. All authors approved the final manuscript.

