## Supplemental Information for "Memorability of Images Positively Influences Associative Memory Retention"

### Supplementary Information for Memorability of Images Positively Influences Associative Memory Retention

#### 1. Supplementary methods

##### 1.1 Stimuli Selection of Experiment 1

Experiment 1 used an image-word associative memory paradigm, where each trial presented an image as the cue and a word as the target. Both cues and targets were selected from the THINGS database[1], which contains 1,854 object concepts, each represented by an average of 14.08 (±2.71) images with established memorability scores. With the exception of 10 concepts, all concepts in the database also have word embeddings, enabling quantitative assessment of semantic similarity between cues and targets.

We calculated the mean (0.790) and standard deviation (0.082) of memorability scores for all images. Images with memorability scores between the mean plus one standard deviation and the mean plus two standard deviations (0.871–0.953) were classified as high memorability (HM), while those between the mean minus two standard deviations and the mean minus one standard deviation (0.626–0.708) were classified as low memorability (LM). In total, 1,515 concepts had at least one HM image and 1,371 had at least one LM image.

Semantic relatedness between all concept pairs was computed as 1 minus the cosine distance of their word embeddings. The mean semantic relatedness was 0.272 (*SD* = 0.106). Pairs with semantic relatedness values in the top 0.5% (0.728–0.975) were classified as high semantic relatedness (HS), while those with values from the minimum up to the mean minus one standard deviation (-0.123–0.166) were classified as low semantic relatedness (LS). In total, 1,699 concept pairs were classified as HS and 239,263 as LS.

Three research assistants removed concepts unfamiliar to Chinese (e.g., "totem pole") from the dataset. This filtering reduced the set to 1,465 concepts. Of these, 1,280 had HM images, 1,143 had LM images, 1,049 concept pairs met the HS criteria, and 150,176 met the LS criteria.

We identified all possible concept pairs classified as high semantic similarity (HS) or low semantic similarity (LS), in which the cue image was either high memorability (HM) or low memorability (LM). The number of eligible pairs for each condition was as follows:

| **Stimulus Condition** | **Number of Pairs** |
|---|---|
| **HM+HS** | 994 |
| **LM+HS** | 963 |
| **HM+LS** | 145,049 |
| **LM+LS** | 140,513 |

We randomly selected 30 concept pairs from each condition. To control for potential confounding factors, we extracted low-level image statistics (color, brightness, contrast, spatial frequency) using the Natural Image Statistical Toolbox[2], and obtained word frequency data from the Peking University CCL corpus[3] as well as word length for each target. We then conducted ANOVAs to confirm that there were no significant differences in image or word features across experimental conditions (implemented in Python v3.11.5 with pandas v1.5.3 and scipy v1.10.1). If any significant differences were detected ($P < 0.05$), a new set of 30 pairs was randomly drawn until all features were matched (all $P > 0.05$). The final set of concept pairs was distributed across the memorability-semantic relatedness space as shown in **Fig. S1**.

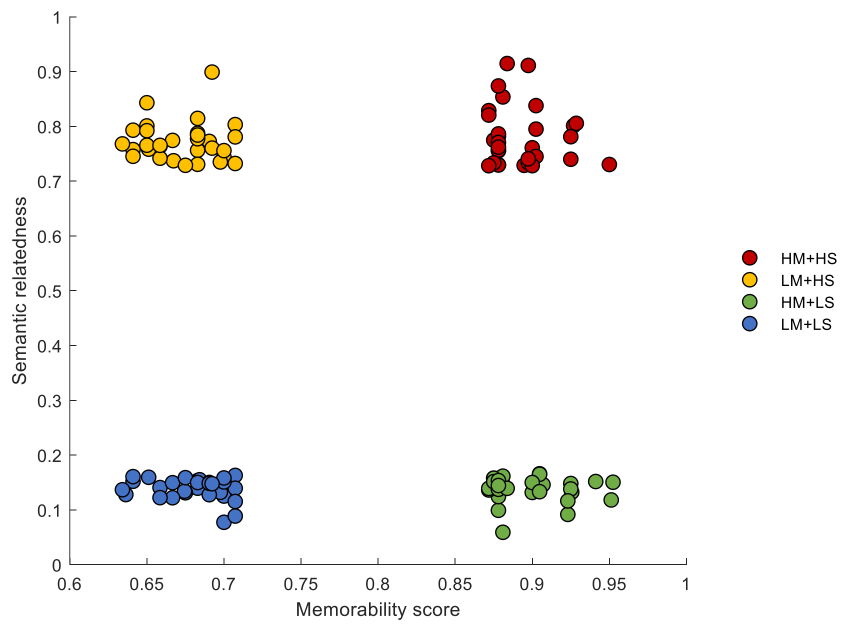


**Figure S1. The distribution of selected concept pairs of Experiment 1 in memorability-sematic**

**relatedness space.**

#### 1.2 Stimuli Selection of Experiment 2

To evaluate the generalizability of our findings, Experiment 2 used a new set of stimuli rather than reusing those from Experiment 1. The procedure for selecting concept pairs was identical to that of Experiment 1. The distribution of these pairs within the memorability-semantic relatedness space is shown in **Fig. S2**.

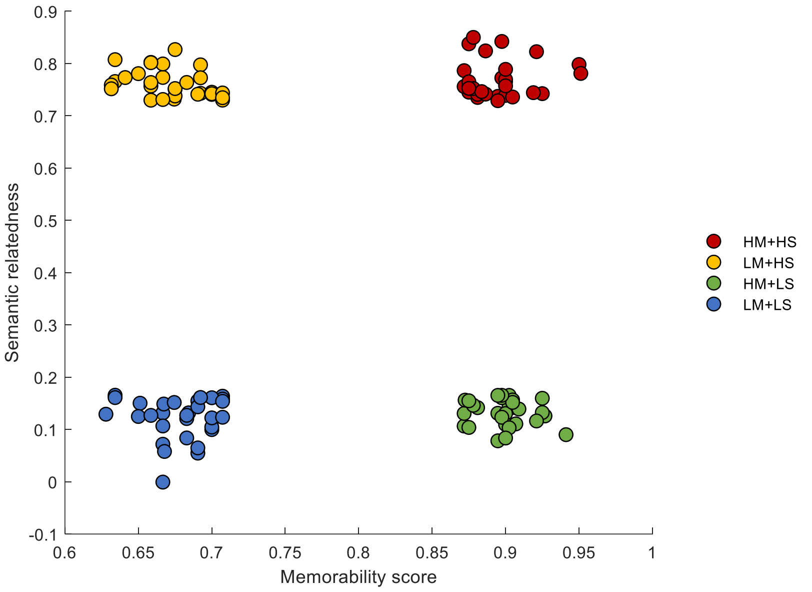


**Figure S2. The distribution of selected concept pairs of Experiment 2 in memorability-sematic relatedness space.**

#### 1.3 Stimuli Selection of Experiment 3

Given that most Chinese college students have some proficiency in English, we selected Korean as the target language to minimize interference from English or related languages. To control for the potential influence of word length on memory, only two-syllable Korean words were included. The images paired with these words were drawn from the THINGS database[1]. We initially identified 1,101 concepts that had both high-memorability (HM) and low-memorability (LM) images available. To avoid carryover effects from Experiment 1–since some participants took part in both studies–we excluded 120 concepts that had appeared previously. After further restricting the set to concepts paired with two-syllable Korean words, 188 eligible concepts remained. Finally, three research assistants reviewed the list to exclude concepts unfamiliar to Chinese participants, resulting in a final set of 187 concepts.

For each concept, we selected one HM image and one LM image, resulting in a paired image set for the two-syllable Korean vocabulary. Sixty concepts were then randomly chosen to serve as stimuli for the fixed-time experiment, with 30 assigned to Group A and the other 30 to Group B. To ensure that potential confounding features–such as image color, brightness, contrast, and spatial frequency, as well as word length and frequency–were balanced across conditions, we extracted low-level image statistics using the Natural Image Statistical Toolbox[2]. We then conducted ANOVAs on the four groups of stimuli (Group A HM, Group B HM, Group A LM, Group B LM) to test for significant differences in image characteristics (implemented in Python [v3.11.5] using pandas [v1.5.3] and scipy [v1.10.1]). If significant differences were detected (any $P < 0.05$), a new random sample of 60 concepts was drawn and the process was repeated until no differences remained (all $P > 0.05$).

For the self-paced experiment, an additional 60 concepts were randomly selected from the remaining pool, excluding those already used in Groups A and B. Thirty concepts were assigned to Group X and the other thirty to Group Y. As with the fixed-time experiment, low-level image statistics (color, brightness, contrast, and spatial frequency) were calculated for each image using the Natural Image Statistical Toolbox[2]. ANOVAs were then conducted to compare image features across the four groups (Group X HM, Group Y HM, Group X LM, Group Y LM), implemented in Python (v3.11.5) using pandas (v1.5.3) and scipy (v1.10.1). If any significant differences were detected ($P < 0.05$), the selection process was repeated until no significant differences remained (all $P > 0.05$).

### 2. Supplementary analyses

#### 2.1 The interactive effects of memorability and semantic relatedness in distinct age groups in Experiment 1

Under the high semantic relatedness (HS) condition, highly memorable (HM) images led to significantly higher recall accuracy than low memorability (LM) images in the young adult group ($F(1, 151) = 6.734$, $P = 0.010$, $\eta^2 = 0.043$; mean difference = 1.747%, 95% CI = [0.417%, 3.077%]). However, no significant effect of image memorability was observed for the middle-aged group ($F(1, 53) = 0.109$, $P = 0.743$, $\eta^2 = 0.002$; mean difference = 0.417%, 95% CI = [-2.120%, 2.954%]) or the older adult group ($F(1, 50) = 1.800$, $P = 0.186$, $\eta^2 = 0.035$; mean difference = -2.160%, 95% CI =

[-5.395%, 1.074%]).

In contrast, under the low semantic relatedness (LS) condition, HM images significantly enhanced recall accuracy compared to LM images across all age groups: young adults ($F(1, 151) = 173.213$, $P < 0.001$, $\eta^2 = 0.534$; mean difference = 11.376%, 95% CI = [9.668%, 13.084%]), middle-aged adults ($F(1, 53) = 68.769$, $P < 0.001$, $\eta^2 = 0.565$; mean difference = 11.152%, 95% CI = [8.455%, 13.850%]), and older adults ($F(1, 50) = 45.502$, $P < 0.001$, $\eta^2 = 0.476$; mean difference = 9.334%, 95% CI = [6.554%, 12.113%]).

#### 2.2 The interaction effects of memorability and other factors in Experiment 2

The interaction between memorability and group was not significant ($F(1, 98) = 2.262$, $P = 0.136$, $\eta^2 = 0.023$), whereas there was a significant interaction between semantic relatedness and group ($F(1, 98) = 5.391$, $P = 0.022$, $\eta^2 = 0.052$), as well as between memorability and semantic relatedness ($F(1, 98) = 34.043$, $P < 0.001$, $\eta^2 = 0.258$). Simple effects analyses indicated that, regardless of semantic relatedness, HM images led to significantly higher recall accuracy than LM images under both high semantic relatedness ($F(1, 99) = 12.649$, $P = 0.001$, $\eta^2 = 0.113$; mean difference = 3.319%, 95% CI = [1.467%, 5.171%]) and low semantic relatedness ($F(1, 99) = 148.235$, $P < 0.001$, $\eta^2 = 0.600$; mean difference = 11.216%, 95% CI = [9.388%, 13.044%]). The three-way interaction was not significant ($F(1, 98) = 0.189$, $P = 0.665$, $\eta^2 = 0.002$).

#### 2.3 Comparison of the Memory Enhancement Effect of HM Images Across Semantic Relatedness Conditions in Experiment 2

We further investigated whether the magnitude of the memory enhancement effect–calculated as the difference in recall rates between HM and LM images–varied between adults with normal cognitive aging and those with cognitive impairment. A 2 (semantic relatedness: high vs. low) × 2 (group: cognitively normal vs. cognitively impaired) mixed-design ANOVA was conducted, with the group as the between-subjects factor.

The main effect of semantic relatedness was significant ($F(1, 98) = 70.533$, $P < 0.001$, $\eta^2 = 0.419$): the recall advantage of HM over LM images was substantially greater under the low semantic relatedness (LS) condition ($M = 37.849\%$, $SE = 3.588\%$) than under the high semantic relatedness (HS) condition ($M = 4.872\%$, $SE = 1.402\%$), with a mean difference of -32.977%, 95% CI = [-40.769%, -25.185%]. There was no significant main effect of group ($F(1, 98) = 0.025$, $P = 0.874$, $\eta^2 = 0.000$), nor a significant interaction between semantic relatedness and group ($F(1, 98) = 0.000$, $P = 0.985$, $\eta^2 = 0.000$). These results suggest that the memory enhancement provided by HM images, compared to LM images, is equally robust in both cognitively normal and cognitively impaired older adults.

#### 2.4 Follow-up Analysis of the Cognitively normal and Cognitively Impaired Groups in Experiment 2

To examine whether forgetting rates varied across conditions and participant groups over time, we conducted separate analyses for the 1-day and 7-day follow-up data in Experiment 2. For the 1-day delay, a 2 (memorability: high vs. low) × 2 (semantic relatedness: high vs. low) × 2 (group: cognitively normal older adults vs. cognitively impaired older adults) mixed-design ANOVA was performed, with the group as a between-subjects factor. The results revealed a significant main effect of memorability ($F(1, 76) = 4.009$, $P = 0.049$, $\eta^2 = 0.050$): HM images were associated with significantly lower forgetting rates ($M = 60.330\%$, $SE = 2.785\%$) than LM images ($M = 64.784\%$, SE = 2.029%), with a mean difference of -4.454% (95% CI = [-8.885%, -0.024%]). There was also a significant main effect of semantic relatedness ($F(1, 76) = 160.429$, $P < 0.001$, $\eta^2 = 0.679$), as forgetting rates were higher for LS pairs ($M = 78.814\%$, $SE = 3.037\%$) than for HS pairs ($M = 46.300\%$, $SE = 1.862\%$), with a mean difference of 32.514% (95% CI = [27.401%, 37.626%]). The main effect of group was not significant ($F(1, 76) = 2.227$, $P = 0.140$, $\eta^2 = 0.028$), and neither were the interactions between memorability and group ($F(1, 76) = 1.861$, $P = 0.177$, $\eta^2 = 0.024$), semantic relatedness and group ($F(1, 76) = 1.017$, $P = 0.316$, $\eta^2 = 0.013$), memorability and semantic relatedness ($F(1, 76) = 0.876$, $P = 0.352$, $\eta^2 = 0.011$), nor the three-way interaction ($F(1, 76) = 0.444$, $P = 0.507$, $\eta^2 = 0.006$).

Further analysis within each group revealed that, among normal cognitive aging adults, there were no significant differences in forgetting rates between HM and LM images under either high semantic relatedness (HS; HM: $M = 41.355\%$, $SE = 2.416\%$; LM: $M = 42.187\%$, $SE = 2.417\%$; $F(1, 44) = 0.136$, $P = 0.714$, $\eta^2 = 0.003$, mean difference = -0.832%, 95% CI = [-5.387%, 3.722%]) or low semantic relatedness (LS; HM: $M = 75.869\%$, $SE = 2.623\%$; LM: $M = 77.876\%$, $SE = 3.847\%$; $F(1, 44) = 0.216$, $P = 0.644$, $\eta^2 = 0.005$, mean difference = -2.007%, 95% CI = [-10.711%, 6.697%]). Likewise, for cognitively impaired participants, no significant differences were observed between HM and LM images under either HS (HM: $M = 50.085\%$, $SE = 3.673\%$; LM: $M = 54.499$, $SE = 3.538\%$; $F(1, 36) = 2.664$, $P = 0.111$, $\eta^2 = 0.069$, mean difference = -4.413%, 95% CI = [-9.897%, 1.070%]) or LS conditions (HM: $M = 75.262\%$, $SE = 9.426\%$; LM: $M = 86.247\%$, $SE = 4.013\%$; $F(1, 32) = 2.234$, $P = 0.145$, $\eta^2 = 0.065$, mean difference = -10.984%, 95% CI = [-25.953%, 3.984%]), as shown in **Fig. S3**.

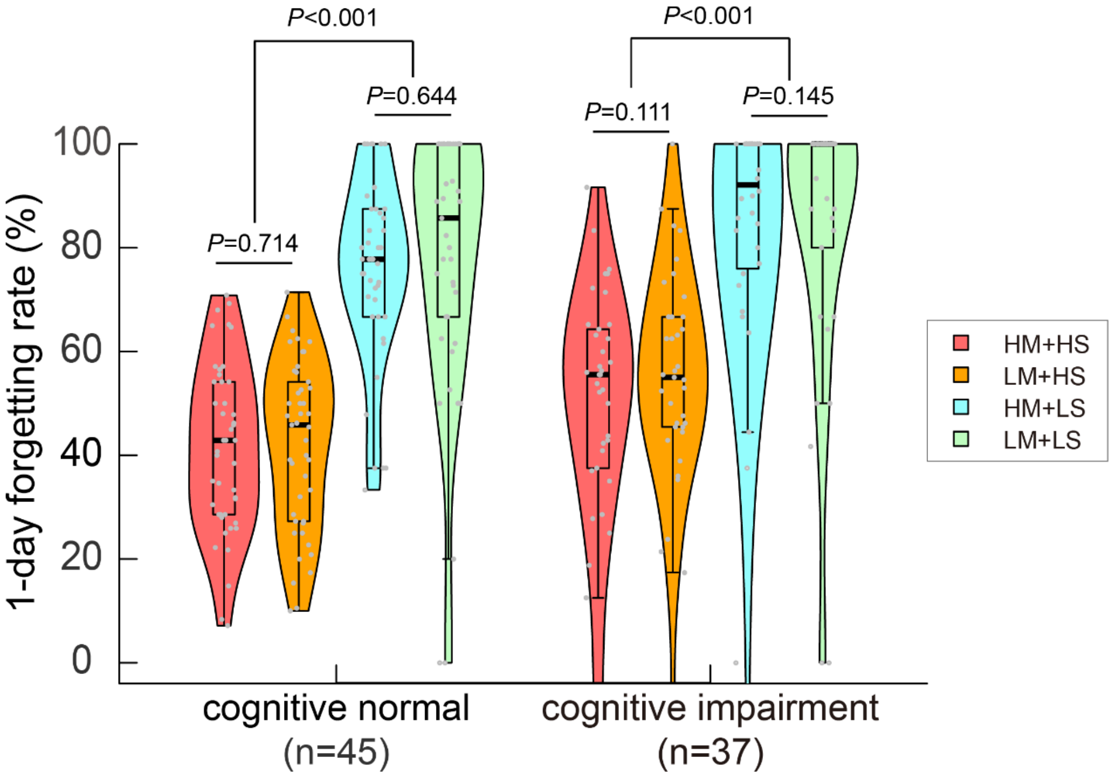


**Figure S3. Forgetting rates under four conditions during the 1-day follow-up test in the normal cognitive aging group and cognitive impairment group.**

At the 7-day follow-up, the mixed-design ANOVA revealed a significant main effect of memorability ($F(1, 72) = 13.144$, $P = 0.001$, $\eta^2 = 0.154$): HM images were associated with significantly lower forgetting rates ($M = 70.101\%$, $SE = 1.882\%$) than LM images ($M = 75.020\%$, $SE = 1.700\%$), with a mean difference of -4.919% (95% CI = [-7.623%, -2.214%]). A robust main effect of semantic relatedness was also observed ($F(1, 72) = 173.095$, $P < 0.001$, $\eta^2 = 0.706$): forgetting was markedly greater for LS pairs ($M = 86.332\%$, $SE = 2.081\%$) than for HS pairs ($M = 58.789\%$, $SE = 1.836\%$; mean difference = -27.543%, 95% CI = [-31.717%, -23.370%]). The main effect of group was significant as well ($F(1, 72) = 8.612$, $P = 0.004$, $\eta^2 = 0.107$), indicating higher forgetting rates in the cognitively impaired group ($M = 77.431\%$, $SE = 2.560\%$) compared to the cognitively normal group ($M = 67.690\%$, $SE = 2.113\%$; mean difference = -9.741%, 95% CI = [-16.358%, -3.124%]). No significant interactions were found between memorability and group ($F(1, 72) = 0.113$, $P = 0.737$, $\eta^2 = 0.002$), semantic relatedness and group ($F(1, 72) = 1.047$, $P = 0.310$, $\eta^2 = 0.014$), nor memorability and semantic relatedness ($F(1, 72) = 0.058$, $P = 0.811$, $\eta^2 = 0.001$), and the three-way interaction ($F(1, 72) = 0.047$, $P = 0.829$, $\eta^2 = 0.001$) was also not significant.

Further within-group analyses revealed that among normal cognitive aging adults, HM images produced significantly lower forgetting rates than LM images under the HS condition ($F(1, 43) = 6.270$, $P = 0.016$, $\eta^2 = 0.127$, mean difference = -5.342%, 95% CI = [-9.645%, -1.040%]). However, under the LS condition, the difference between HM and LM was not significant ($F(1, 43) = 2.573$, $P = 0.116$, $\eta^2 = 0.056$, mean difference = -5.408%, 95% CI = [-12.207%, 1.391%]). For the cognitively impaired group, no significant difference was observed between HM and LM images in the HS condition ($F(1, 33) = 0.948$, $P = 0.337$, $\eta^2 = 0.028$, mean difference = -2.585%, 95% CI = [-7.987%, 2.817%]). In contrast, under the LS condition, HM images led to significantly lower forgetting rates compared to LM images ($F(1, 29) = 7.959$, $P = 0.009$, $\eta^2 = 0.215$, mean difference = -5.102%, 95% CI = [-8.800%, -1.403%]), as shown in **Fig. S4**.

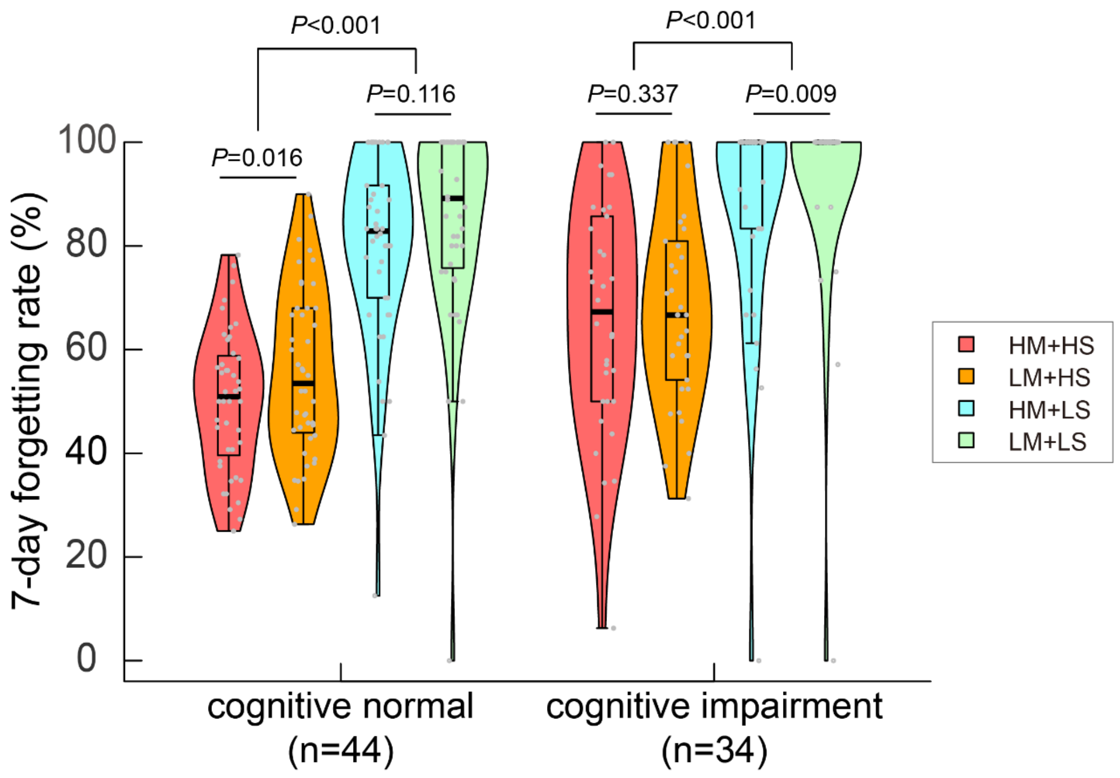


**Figure S4. Forgetting rates under four conditions during the 7-day follow-up test in the normal cognitive aging group and cognitive impairment group.**
